# Antimicrobial Agent Triclosan Inhibits Acetylcholinesterase Activity *in Vitro* in a Dose-Dependent Manner

**DOI:** 10.1101/2021.06.11.448059

**Authors:** Narasimha Pullaguri, R Andrea Kagoo, Anamika Bhargava

## Abstract

The antimicrobial agent, Triclosan, is widely used in many consumer products. It has been designated as a “contaminant of emerging concern (CEC)” because its exposure is known to cause adverse ecological and human health effects. Triclosan is not labelled as GRAS/GRAE (generally recognized as safe and effective), but its use is still prevailing. *In vivo* studies have revealed that exposure to triclosan results in a decreased acetylcholinesterase (AChE) activity. However mechanistic insights into AChE inhibition by triclosan are missing. Using *in vitro* AChE activity assay with purified AChE, we show that triclosan acts as a direct inhibitor of AChE and inhibits AChE activity in a dose-dependent manner. Given the function of AChE, any alteration in its activity can be neurotoxic. Our results provide important mechanistic insights into triclosan induced neurotoxicity with AChE as a target.

## 1. INTRODUCTION

5-chloro-2-(2,4-dichlorophenoxy) phenol, commonly known as Triclosan is a broad-spectrum, synthetic antimicrobial agent that is used in personal care products like toothpastes, soaps, liquid sanitizers, deodorants, mouthwash, shaving gels, beauty care products like lip gloss, moisturizers, children’s toys, kitchenware like cutting boards, semi-automated slicer and other products such as humidifier, vacuum cleaner, interior paints, etc. [1]. High demand for triclosan has resulted in its build-up in drinking and wastewater sources, making it an emerging environmental pollutant [2]. In 2016, triclosan was partially banned by the USA Food and Drug Administration (FDA). However, triclosan still remains in use in rest of the world with its use ever growing [3]. Triclosan is easily absorbed by the skin and is found in various human tissues and fluids. For detailed information about the health effects of triclosan please see review by Weatherly et al [1]. Briefly, it is reported that triclosan affects immune responses, cardiovascular functions, and ROS production [1]. Studies carried out in mice indicate that prenatal exposure to triclosan can induce neurodevelopmental disorders [4]. Recently, we showed that triclosan has the potential to induce neurotoxicity at sublethal concentrations by inhibiting acetylcholinesterase (AChE) enzyme *in vivo* [5]. AChE is a cholinergic enzyme that is found primarily in neuromuscular junctions. It hydrolyses the neurotransmitter acetylcholine thereby precisely controlling the amount of acetylcholine available for neurotransmission at the neuromuscular synapses. Notably, AChE activity is also one of the most common biomarkers of neurotoxicity [6]. Mechanistically, using docking studies, we showed that triclosan can form a stable interaction with the binding pocket of AChE and can compete with the natural neurotransmitter, acetylcholine for binding to the active site of AChE [5]. However, direct inhibition of AChE by triclosan remains to be experimentally confirmed. In the present study, we aimed to determine if triclosan can directly inhibit AChE activity *in vitro*. This has never been shown before. Our results indicate that indeed triclosan can inhibit AChE activity *in vitro* in a dose-dependent manner demonstrating a direct inhibition of AChE by triclosan supporting the *in vivo* results previously observed.

## 2. MATERIALS AND METHODS

### 2.1. Reagents

Triclosan (CAS No. 3380-34-5) and acetone (CAS No. 15168) were purchased from Sigma-Aldrich Co and SR Laboratories, India respectively. For each set of experiments, a fresh stock triclosan solution (4 mg/mL) was made in acetone and stored at 4°C. Different concentrations of triclosan working solutions (with final concentration of triclosan as 0.3 mg/L, 0.6 mg/L and 6 mg/L) were prepared by diluting the triclosan stock solution in 20 mM Tris-HCl buffer (pH 7.5). Since triclosan is dissolved in acetone, therefore a solvent control was included in our experiments where solvent control contained 0.0147 % and 0.147 % final acetone concentration in the triclosan working solutions of 0.6 mg/L and 6 mg/L triclosan, respectively. Neostigmine bromide, a known AChE inhibitor was used as positive control. Neostigmine bromide (CAS No. 114-80-7), AChE from electric eel (type VI-S; 288 U/mg protein) (CAS No. 9000-81-1), and Glutathione (CAS No. 70-18-8) were purchased from Sigma Aldrich Co. 5,5’-dithiobis-(2-nitrobenzoic acid) or Ellman’s reagent (DTNB) (CAS No. 69-78-3), Acetylthiocholine iodide (CAS No. 1886-15-5), Hydrochloric acid (CAS No. 7647-01-0), and Tris(hydroxymethyl) aminomethane (CAS No. 77-86-1) were purchased from HiMedia Laboratories. AChE stock solution (1 mg/ml) was prepared in 20 mM Tris-HCl buffer (pH 7.5) and stored at -20°C. AChE working solution (0.1 µg/ml) was prepared freshly each time in the same buffer for each reaction. Neostigmine Bromide stock solution (20 mg/ml) and working solutions were also prepared in the same buffer. The buffer was filtered through sterile syringe filter (pore size: 0.22 μm) before use.

### 2.2. Measuring Acetylcholinesterase activity *in vitro*

AChE activity was determined using a modified protocol of the Ellman method [7]. It is a spectrophotometric assay that measures the hydrolysis of a thio analog of acetylcholine. The esterase hydrolyses the acetylthiocholine substrate, producing thiocholine, which in turn combines with DTNB [5,5’-dithiobis-(2-nitrobenzorc acid)], forming a yellow-coloured product which can be measured at 412 nm. Glutathione was used as a reference standard because it supplies sulfhydryl groups. 1 µl of 1 mM glutathione solution contains 1 nmol of sulfhydryl groups [8]. Hence, a glutathione standard curve was plotted and used for determination of AChE activity. In some of the inhibition experiments (as indicated against the experiments) the enzyme solution was preincubated with the inhibitor (triclosan or neostigmine bromide) or Tris-HCl buffer in case of controls. Preincubation of enzyme and inhibitor for a predefined period effectively establishes a full equilibrium between the enzyme and the inhibitor [9]. After preincubation of the enzyme with the inhibitor, substrate was added to initiate the enzyme-catalysed reaction.

The blank to control for the non-enzymatic hydrolysis of acetylthiocholine contained a mixture of 20 µl of 0.3 mM DTNB solution (in Tris-HCl buffer), 20 µl of 1.6 mM ATCI (in Tris-HCl buffer) and 215 µl of Tris-HCl buffer. In the reaction wells, 30 µl of Tris-HCl buffer was replaced by AChE solution (0.1 µg/ml). For reactions involving inhibitors, 15 µl of Tris-HCl buffer was replaced by the inhibitor (either triclosan or neostigmine bromide). The blank to control for the non-enzymatic hydrolysis of acetylthiocholine, in the presence of an inhibitor, contained a mixture of 20 µl of 0.3 mM DTNB solution (in Tris-HCl buffer), 20 µl of 1.6 mM ATCI (in Tris-HCl buffer), 15 µl inhibitor solution and 200 µl of Tris-HCl buffer. Absorbance readings were taken at 412 nm using multimode plate reader (Model: Enspire ®, Perkin Elmer). The reaction was monitored for 20 min and the absorbance was noted every minute. In most of the experiments the saturation in enzyme activity was reached in 10 min and therefore activity is reported at 10 min.

### 2.3. Analysis and Statistics

All experiments were carried out in technical triplicates in 96-well plates with a minimum of two trials and a maximum of six trials. The data represents mean ± SEM for the given number of experiments. Percent inhibition was calculated as 100 - ((concentration of product formed in 10 minutes in case of inhibitory compound/concentration of product formed in 10 min in case of control) × 100). Graphpad Prism 7 was used for analysis. One sample t-test against a hypothetical value of zero was performed to check whether percent inhibition was significantly different from control or not. *P* < 0.05 was considered significant. * denotes *P* ≤ 0.05 and *** denotes *P* ≤ 0.005.

## 3. RESULTS AND DISCUSSION

### 3.1. Acetylcholinesterase activity *in vitro*

First, we optimised the AChE enzyme and substrate concentration required to obtain measurable AChE activity in our lab conditions. We tested several substrate and enzyme concentrations from the published literature. We obtained reproducible results at the substrate concentration of 0.125 mM for two different enzyme concentrations as shown in figure 1. In the figure 1, enzyme activity is plotted as nmoles of product obtained after 10 minutes of the reaction at two different enzyme concentrations. From these experiments, we chose the enzyme concentration of 0.06 μg/ml for further experiments.

**Figure 1.**
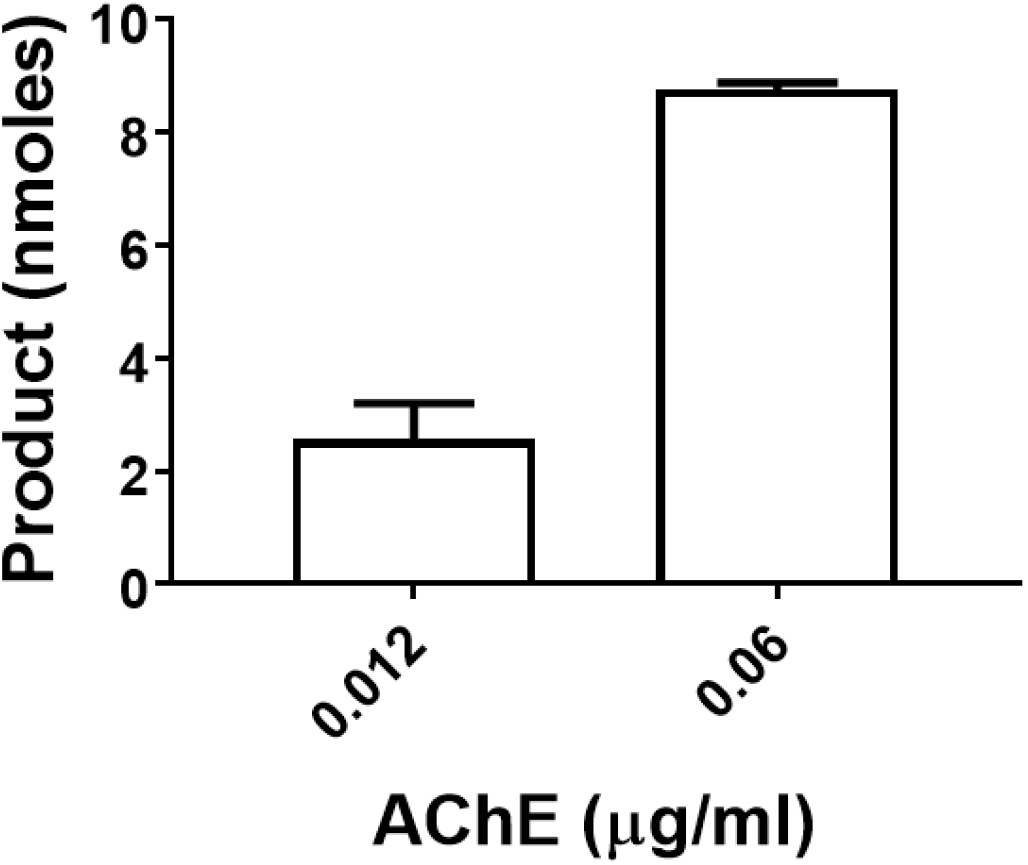
Acetylcholinesterase activity at two different enzyme concentrations (0.012 μg/ml and 0.06 μg/ml). Enzyme activity is plotted as nmoles of product obtained after 10 minutes of the reaction. Final concentration of the substrate was 0.125 mM. The data represents Mean ± SEM of 3 individual experiments with 3 technical replicates each.

### 3.2. Inhibition of acetylcholinesterase activity by triclosan

In order to observe the inhibition of AChE activity, we first used triclosan at concentrations 0.3 and 0.6 mg/L as used previously *in vivo* [5]. Figure 2 shows AChE activity inhibition by triclosan at the two tested concentrations. We also performed AChE activity assay with a known inhibitor, neostigmine bromide as a positive control [10]. As expected, we observed a dose-dependent inhibition of AChE activity by neostigmine bromide (Figure 2), however no significant reduction in the AChE activity by triclosan was observed (Figure 2). Dose-dependent inhibition of AChE activity by neostigmine bromide at four different concentrations is also shown in the supplementary figure 1. These four different concentrations of neostigmine bromide were chosen based on the IC_50_ of neostigmine bromide [11].

**Figure 2.**
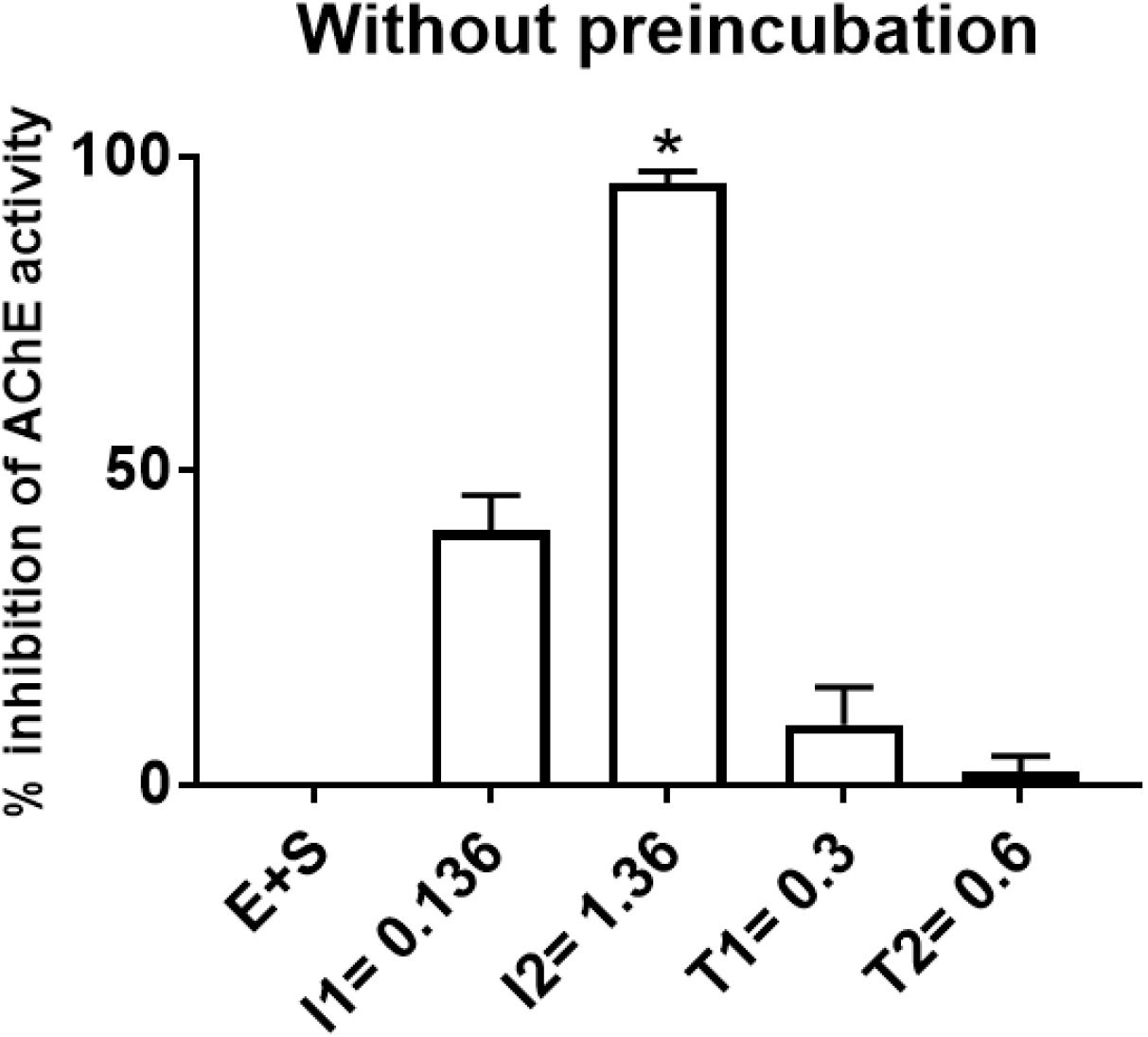
Inhibition of acetylcholinesterase activity by triclosan without preincubation. Percentage inhibition observed in case of control (E+S), Neostigmine bromide (E+S+I1), Neostigmine Bromide (E+S+I2), Triclosan (E+S+T1) and Triclosan (E+S+T2) after 10 minutes of reaction. Final concentration of substrate and AChE was 0.125 mM and 0.06 μg/ml respectively. One sample t-test against a hypothetical value of zero was performed. * *P* ≤ 0.05. E-Enzyme, S-Substrate, I1-Neostigmine bromide inhibitor (0.136 μM), I2-Neostigmine bromide inhibitor (1.36 μM), T1-Triclosan (0.3 mg/L), T2-Triclosan (0.6 mg/L). The data represents Mean ± SEM of 2 individual experiments with 3 technical replicates each.

Please note that in the above experiments, triclosan or neostigmine bromide were added after the substrate addition and if triclosan was a weak inhibitor it may not have been able to compete with the substrate. Therefore, we performed the next set of reactions with preincubation of the enzyme with the triclosan followed by addition of the substrate. For this, we first standardised the preincubation time by preincubating the enzyme with buffer for different times (Supplementary figure 2). We observed that AChE activity was not lost or reduced even after 30 minutes of preincubation with the assay buffer-Tris-HCl (Supplementary figure 2). Hence, 30 min preincubation was considered optimum and used in our next experiments. We further increased triclosan concentration and reduced enzyme concentration to eliminate no binding due to low triclosan concentration. Figure 3 shows the percentage inhibition of AChE activity by triclosan and neostigmine bromide using preincubation protocol. Neostigmine bromide (0.136 µM), used as the positive control, showed a percentage inhibition of 91.45 ± 1.48. The lower concentration of triclosan i.e., 0.6 mg/L showed a percentage inhibition of 16.63 ± 4.66 while the higher concentration of triclosan i.e., 6 mg/L showed a percentage inhibition of 38.47 ± 3.92. This indicates that indeed triclosan could inhibit AChE activity *in vitro*, however a preincubation of AChE with triclosan was required to observe the inhibition at our tested concentrations.

**Figure 3:**
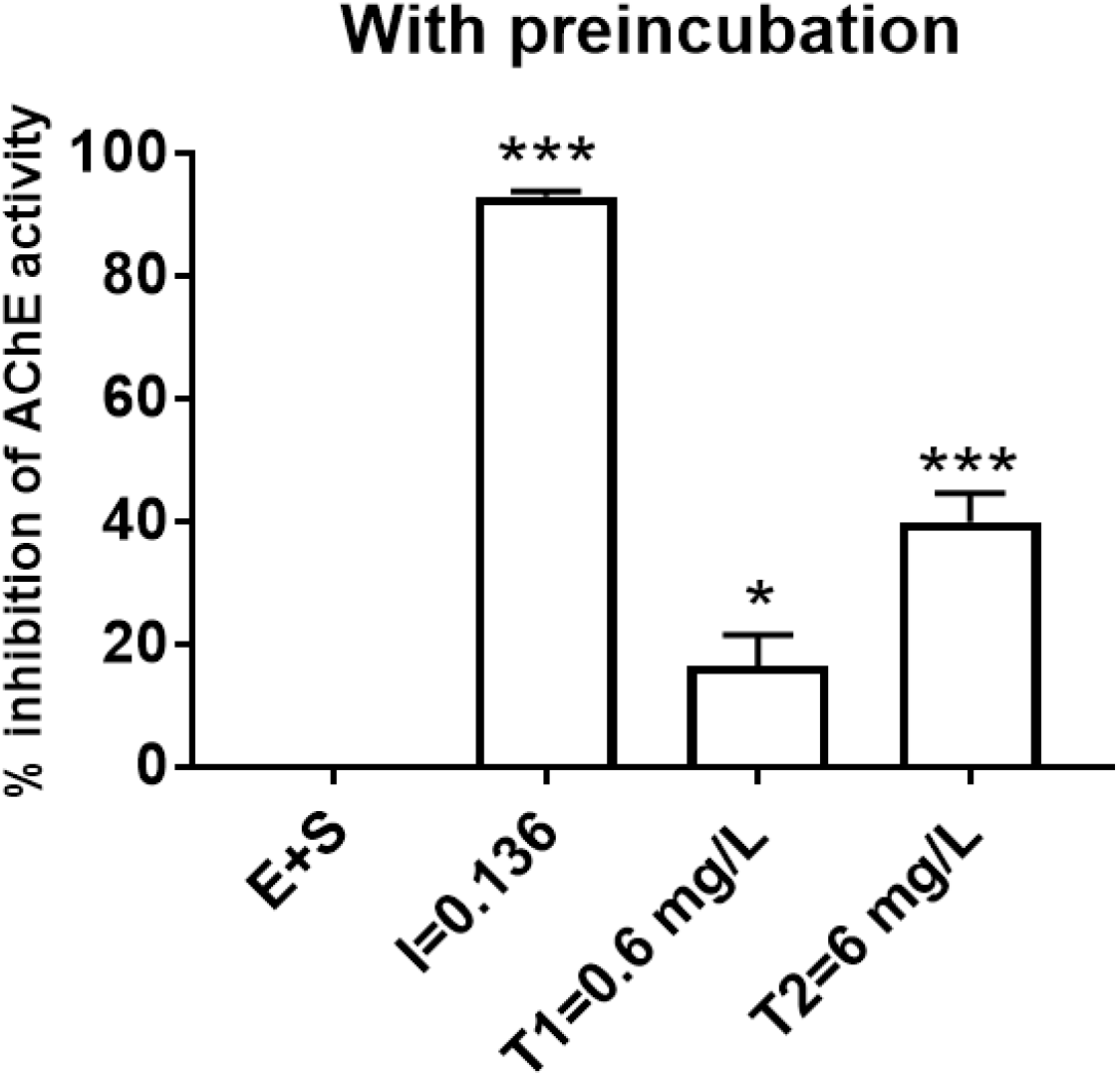
Inhibition of acetylcholinesterase activity by triclosan with preincubation. Percentage inhibition observed in case of control (E+S), Neostigmine bromide (E+S+I), Triclosan (E+S+T1) and Triclosan (E+S+T2) after 10 minutes of the reaction. Final concentration of substrate and AChE was 0.125 mM and 0.012 μg/ml respectively. One sample t-test against a hypothetical value of zero was performed. * *P* ≤ 0.05 and *** *P* ≤ 0.005. E-Enzyme, S-Substrate, I-Neostigmine bromide inhibitor (0.136 μM), T1-Triclosan (0.6 mg/L), T2-Triclosan (6 mg/L). The data represents Mean ± SEM of 6 individual experiments with 3 technical replicates each.

Overall, our results indicate that triclosan can indeed inhibit AChE directly, although at a lower level than the existing established inhibitors. However, it may be noted that, *in vivo*, any alteration in the AChE activity may lead to altered amount of acetylcholine available at the neuromuscular synaptic junctions and thus will impact neurotransmission. Our results are in agreement with *in vivo* studies showing triclosan inhibition of AChE and provide important mechanistic details to support the *in vivo* results. Our results also support the fact that triclosan can be neurotoxic and therefore its use must be regulated.

## 4. CONCLUSION

In this study we show direct evidence of triclosan inhibition of acetylcholinesterase using *in vitro* enzymatic assay. Our results support the *in vivo* data and provide important mechanistic understanding of triclosan induced neurotoxicity. We also believe that our results provide important scientific data in support of restricting or regulating the use of triclosan in future.

## Author contributions

**NP**: Idea for the article, Experiments, Analysis, Literature search, Drafting, Reviewing and Editing **AK**: Experiments, Analysis, Illustrations, Literature search, Drafting, Reviewing and Editing; and **AB**: Idea for the article, Analysis, Drafting, Reviewing and Editing, Illustrations.

## Acknowledgements

This work was supported by the Indian Institute of Technology Hyderabad (IITH) and ECR-SERB-DST (ECR/2017/000242) grants to AB and Ministry of Education, India fellowship to NP and AK.

## Supplementary figures

**Supplementary Figure 1.**
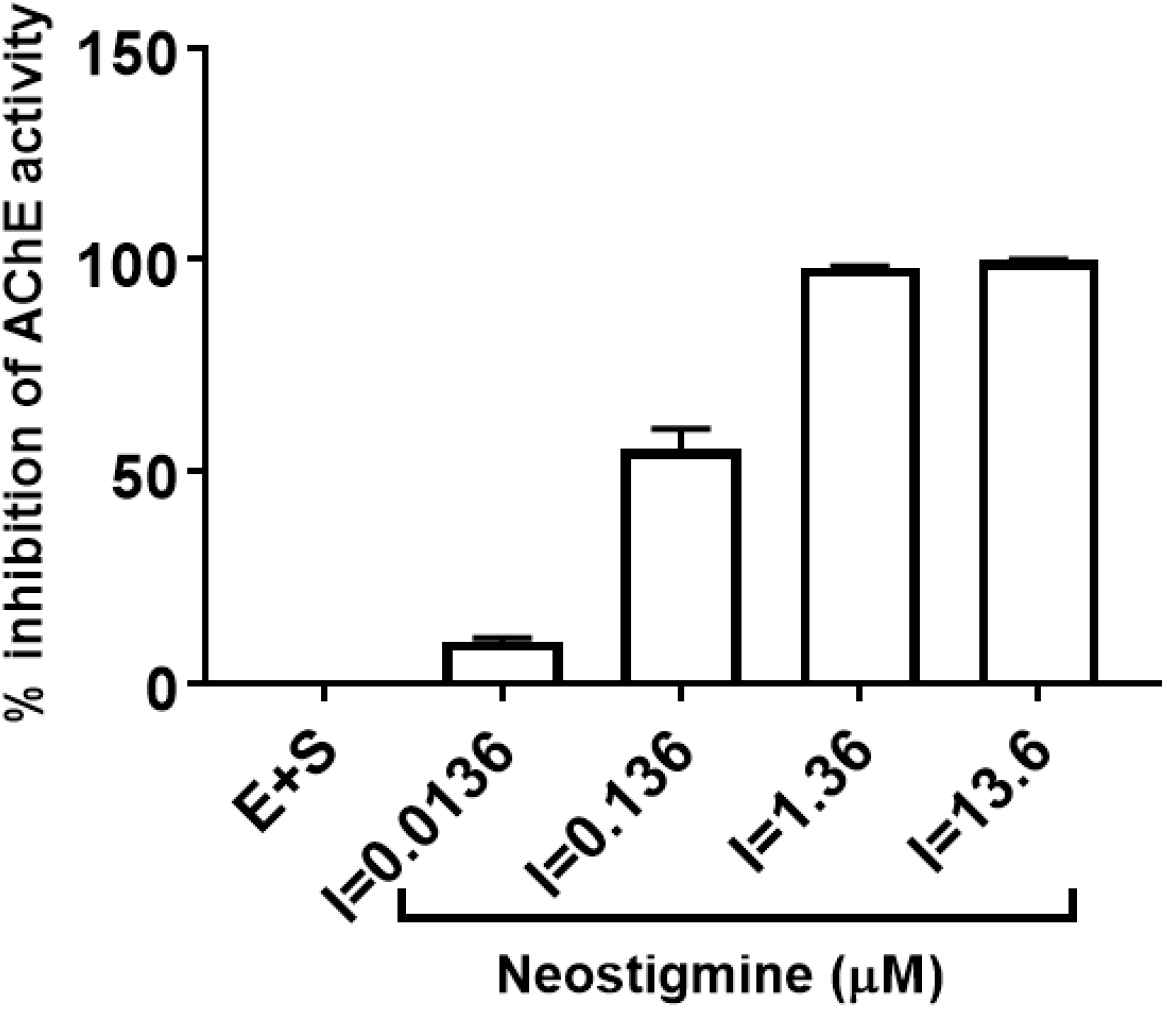
Inhibition of AChE by Neostigmine bromide. Percentage inhibition observed in case of control (E+S) and Neostigmine bromide at various concentrations as indicated after 10 minutes of the reaction. Final concentration of substrate and AChE was 0.125 mM and 0.06 μg/ml respectively. E-Enzyme, S-Substrate, I-Neostigmine bromide inhibitor. The data represents Mean ± SEM of 2 individual experiments with 3 technical replicates each.

**Supplementary Figure 2.**
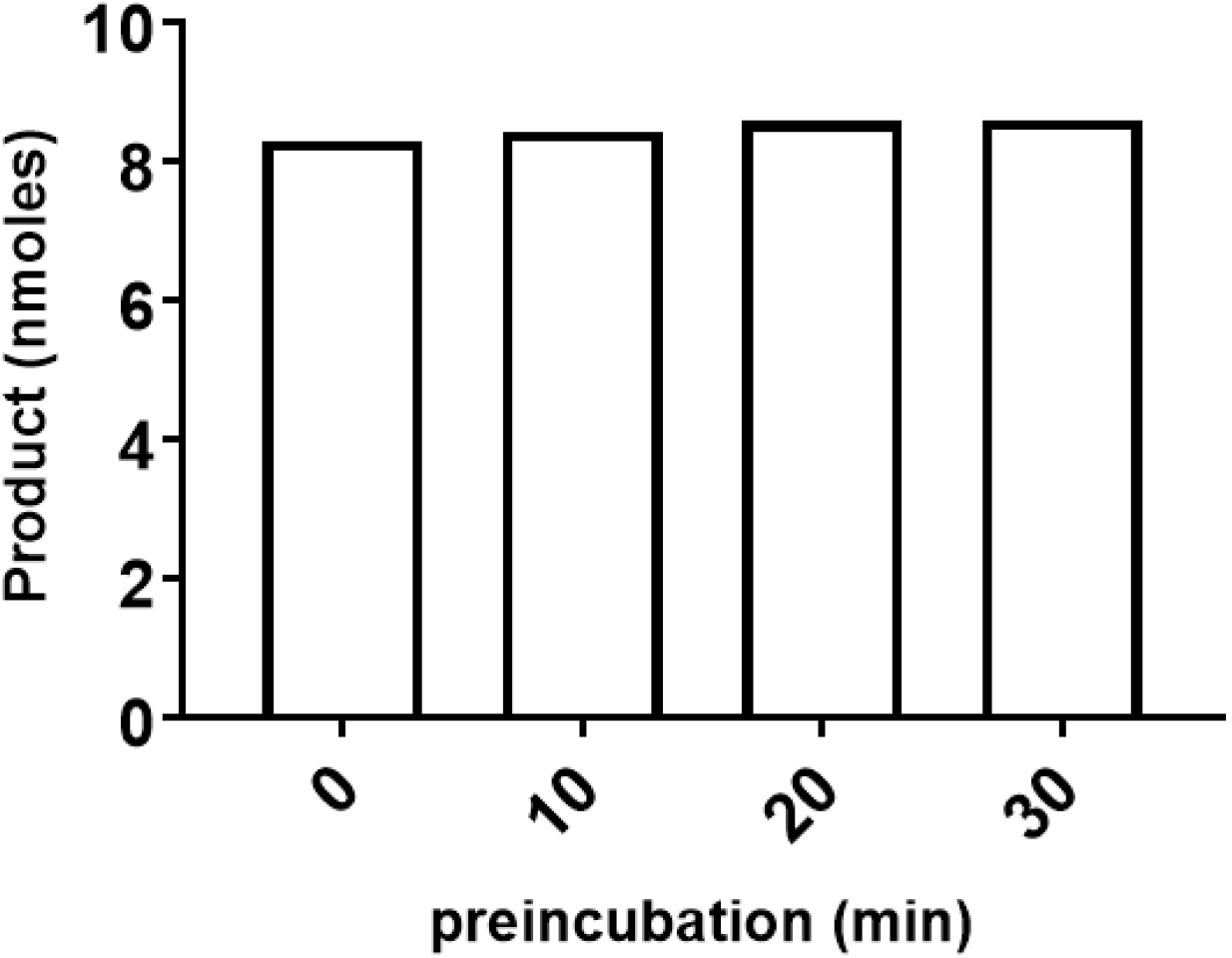
Acetylcholinesterase activity using variable preincubation time. During preincubation, the enzyme was incubated only with the buffer. Afterwards, substrate was added. Enzyme activity is plotted as nmoles of product obtained after 10 minutes of the reaction. Final concentration of the substrate and AChE was 0.125 mM and 0.06 μg/ml respectively. Readings were taken for 10 min. The experiment was performed with 3 technical replicates.

